# Galectin-3 binds to the RGD-binding site in a glycan-independent manner and to the allosteric site and activates integrins αvβ3, αIIbβ3, and α5β1

**DOI:** 10.64898/2026.02.05.704096

**Authors:** Yoko K. Takada, Yu-Jui Yvonne Wan, Yoshikazu Takada

## Abstract

Galectin-3 (Gal3) is one of the most pro-inflammatory proteins and a biomarker of inflammatory diseases and cancer. Previous studies showed that Gal3 binds to αv and β1 integrins but it is unclear how Gal3 binds to integrins. Here, we show that Gal3 bound to soluble αvβ3 and αIIbβ3 integrins in 1 mM Mn^2+^ in cell-free conditions in a glycan-independent manner. Docking simulation predicts that Gal3 binds to the classical RGD-binding site (site 1) of αvβ3, but the predicted Gal3-binding site does not include galactose-binding site. RGDfV or eptifibatide inhibited Gal3 binding to αvβ3 and αIIbβ3, respectively, but lactose, pan-galectin inhibitor, did not inhibit Gal3 binding to integrins. Point mutations of the predicted site 1 binding interface of Gal3 effectively inhibited Gal3 binding to site 1. Site 2 is involved in pro-inflammatory signaling (e.g., TNF and IL-6 secretion) and we previously showed that pro-inflammatory cytokines (e.g., CCL5 and TNF) bind to site 2 and allosteric integrin activation. Docking simulation predicts that Gal3 binds to site 2 of αvβ3 and α5β1. We found that Gal3 induced allosteric activation of soluble integrins αvβ3, αIIbβ3, and α5β1 in 1 mM Ca^2+^ in cell-free conditions. Point mutations in the predicted site 2-binding interface inhibited Gal3-induced integrin activation, suggesting that Gal3 binding to site 2 is required for Gal3-induced integrin activation. Known anti-inflammatory agents, Ivermectin, NRG1, and FGF1 inhibited integrin activation induced by Gal3 in αvβ3 and αIIbβ3. These findings suggest that Gal3 binding to site 2 may be a potential mechanism of pro-inflammatory and pro-thrombotic action of Gal3.

## Introduction

Integrins are a superfamily of alpha-beta heterodimers of cell adhesion receptors that recognize extracellular matrix (ECM) proteins, cell-surface molecules (e.g., ICAM-1 and VCAM-1), and small proteins (e.g., growth factors such as NRG1, CCL5, FGF2) (3).

We performed virtual screening to protein data bank (PDB) with the headpiece of integrin αvβ3 as a target using docking simulation. Interestingly, we discovered that multiple pro-inflammathetory proteins, including cytokines (e.g., FGF1, FGF2, IGF1, IGF2, CD40L, VEGF165, CX3CL1, CXCL12, CCL5), pro-inflammatory proteins such as sPLA2-IIA, CD62P bind to the classical ligand-binding site (site 1).

The integrin ligands are known to bind to the classical RGD-binding site in the headpiece (site 1). Integrins are involved in growth factor signaling (integrin-growth factor crosstalk) since antagonists to integrins suppress growth factor signaling (4-6). Previous studies showed that several growth factors induce integrin-growth factor-cognate growth factor ternary complex on the surface. Growth factor mutants defective in integrin binding were defective in signaling functions and act as antagonists of growth factor signaling. These findings suggest that the ternary complex formation is critical for growth factor signaling (Ternary complex model).

Notably, we found that several pro-inflammatory cytokines or pro-inflammatory proteins (e.g., FGF2, CX3CL1, CXCL12, CCL5, CD40L, and sPLA2-IIA) bind to the allosteric site (site 2), which is distinct from site 1, and activate integrins in an allosteric manner (ref). Site 2-derived peptides from integrin b subunits bound to these ligands and suppressed allosteric activation by these ligands. These findings suggest that the binding to these ligands to site 2 is required for allosteric activation of integrins. It has been reported that 25-hydroxycholesterol (25HC), a major inflammatory mediator, binds to site 2 and activates integrins and induces inflammatory signals (e.g., secretion of IL-6 and TNF) in macrophages (7). Notably, we found that ivermectin, known anti-parasitic and anti-inflammatory small molecule, binds to site 2 and inhibits integrin activation by pro-inflammatory cytokines (8), suggesting that ivermectin is an antagonist of site 2-mediated inflammatory signaling. Also, known anti-inflammatory cytokines, NRG1 and FGF1, bind to site 2 and inhibit integrin activation by multiple pro-inflammatory cytokines (2, 9). We propose that site 2 is involved in inflammatory signaling.

Galectin-3 (Gal3) is a galactoside-binding lectin which is important in numerous biological activities in various organs, including cell proliferation, apoptotic regulation, inflammation, fibrosis, and host defense (10, 11). Gal3 is predominantly located in the cytoplasm and expressed on the cell surface, and then often secreted into biological fluids, like serum and urine. It is also released from injured cells and inflammatory cells under various pathological conditions. Many studies have revealed that Gal3 plays an important role as a diagnostic or prognostic biomarker for certain types of heart disease, kidney disease, viral infection, autoimmune disease, neurodegenerative disorders, and tumor formation. Gal3 positively or negatively affects cancer progression and metastasis (12-14). The mechanism of pro-inflammatory action of Gal3 is unclear. Gal3 has been reported to interact with β1 integrins (α2β1, α3β1). Also, Gal3 enhances cell adhesion to ECM (15), binds to αv integrins (16) and α5β1 integrin (17). We hypothesized that Gal3 activates integrins and induces pro-inflammatory signals by binding to site 2, like pro-inflammatory cytokines.

It is unclear how Gal3 binds to integrins. It has been proposed that Gal3 binding to integrins is mediated by carbohydrate binding. We previously discovered that another carbohydrate-binding P-selectin (CD62P) lectin domain binds to integrins through protein-protein interaction (18). P-selectin lectin domain binds to site 2 and activated integrins through binding to site 2. The present studied if Gal3 binds to site 1 of integrins in a glycan-independent manner and RGD-dependent manner. We also studied if Gal3 binds to site 2 and allosterically activates integrins as several pro-inflammatory cytokines.

## Results

### Gal3 binds to soluble αvβ3 and αIIbβ3 in 1 mM Mn^**2+**^

Binding of Gal3 to integrins was studied in ELISA-type binding assay using soluble in 1 mM Mn^2+^, which fully activate integrins. We found that αvβ3 and αIIbβ3 bound to immobilized Gal3 in a dose-dependent manner (Figure 1). RGDfV is a specific inhibitor of αvβ3. We found that the binding of soluble αvβ3 to immobilized Gal3 in 1 mM Mn^2+^ was inhibited by RGDfV (1 mM). Eptifibatide, a specific inhibitor for αIIbβ3, inhibited Gal3 binding to αIIbβ3 (0.65 µg/ml). These findings suggest that Gal3 binds to the classical RGD-binding site (site 1) of αvβ3 and αIIbβ3 in an RGD-dependent manner. Lactose did not significantly affect Gal3 binding to αvβ3 and αIIbβ3 (up to 50 mM), which is consistent with the idea that Gal3 bind to integrins in a protein-protein interaction.

**Figure 1.**
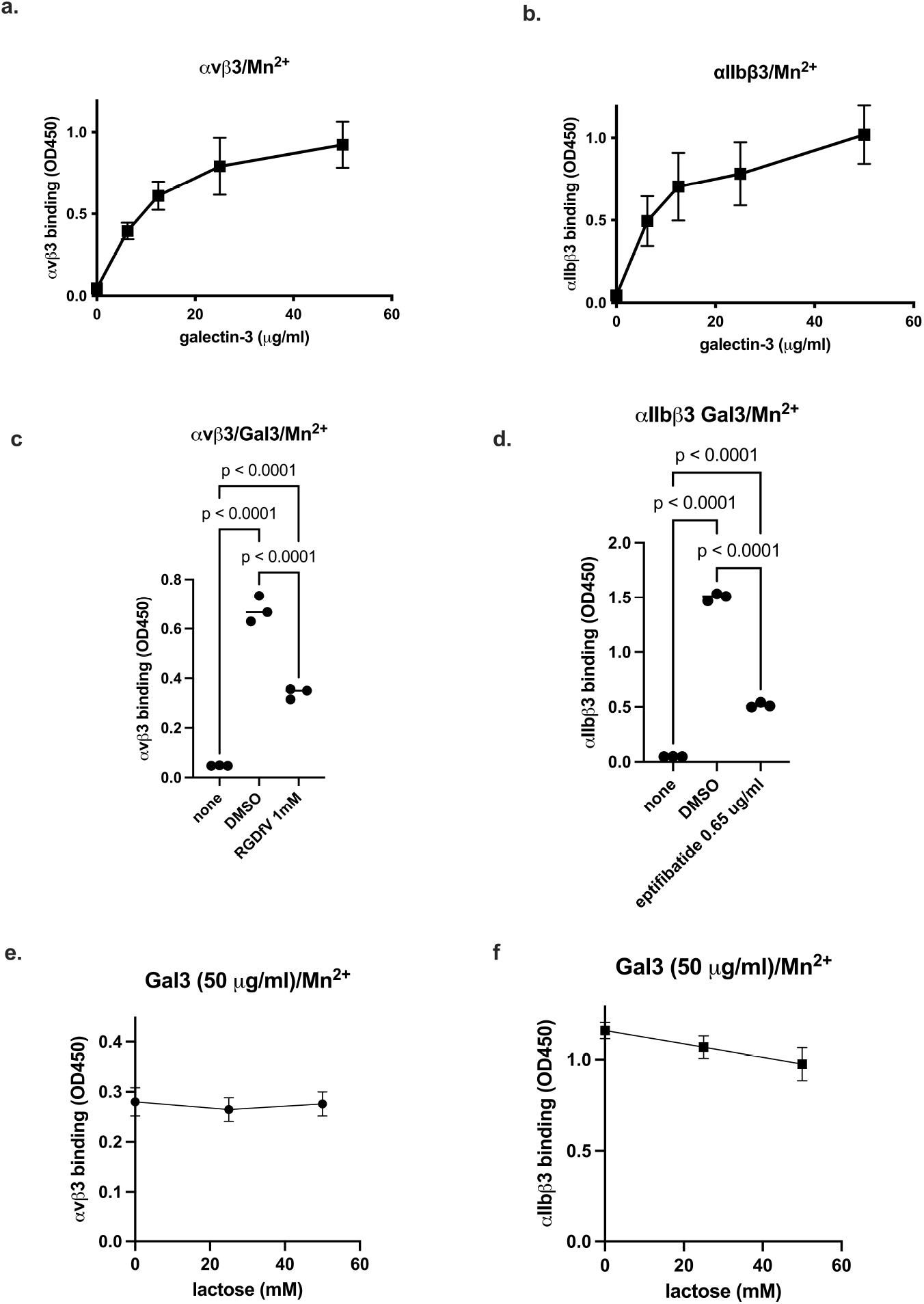
Gal3 binding to soluble integrins αvβ3 and αIIbβ3 in 1 mM Mn^2+^. (a)(b) Gal3 binding to soluble integrins was measured as described in Methods. Briefly, Gal3 was immobilized to wells and incubated with soluble αvβ3 or αIIbβ3. Bound integrins were measured using anti-β3 and HRP-conjugated anti mouse IgG. Data is shown as means +/-SD of triplicate experiments. (c)(d) Effect of integrin inhibitors RGDfV (1 mM) and eptifibatide (0.65 µg/ml) on Gal3 binding to soluble αvβ3 and αIIbβ3, respectively, was measured. (e)(f) Effect of lactose (up to 50 mM) on Gal3 binding to to soluble αvβ3 and αIIbβ3, respectively, was measured. There was no significant effect of lactose on Gal3 binding to integrins. Data is shown as means +/-SD of triplicate experiments.

### Docking simulation predicts that Gal3 binds to the classical RGD-binding site (site 1) of integrin αvβ3

We performed docking simulation between Gal3 (1A3K.pdb) and integrin αvβ3 headpiece (open headpiece conformation, 1L5G.pdb). The 3D structures of open and closed-headpiece forms of integrin were well defined in αvβ3 (1L5G and 1JV2.pdb, respectively). Therefore, we showed docking models of only αvβ3 are shown. Carbohydrate molecules were removed from the αvβ3 headpiece or Gal3. The simulation predicts that Gal3 binds to the classical ligand (RGD) binding site (site 1) of αvβ3 (docking poses are clustered at docking energy 23 kcal/mol) (Figure 2). Predicted amino acid residues involved in Gal3-αvβ3 interaction are shown in Table 1.

**Table 1.** Predicted amino acid residues that are involved in Gal3-integrin αvβ3 binding.

| 1A3K (-23) | $\alpha v$ | $\beta 3$ (1L5G) |
| --- | --- | --- |
| **Arg151**, Asn153, Asp154, Asn166, Asn174, Thr175, **Lys176**, Leu177, Asp178, Asn179, Asn180, Trp181, Gly182, Arg183, Glu184, Glu185, Arg186, **Arg224**, **Lys226**, Lys227, Glu230 | Ala149, Asp150, Tyr178, Trp179, Ala215, Ile216, Asp218, Asp219, Aeg248, | Tyr166, Asp179, Met180, Lys181, Thr182, Arg214, Asn215, Arg216, Asp217, Ala218, Pro219, Glu220, Asp251, Ala252, Lys253, Thr311, Glu312, Asn313, Val314, Asn316, Leu333, Ser334, Met335, Asp336, Ser337 |

Amino acid residues within 0.6 nm between galectin-3 and $\alpha\beta 3$ were selected using Pdb Viewer (version 4.1). Amino acid residues of Gal3 selected for mutation are shown in bold.

**Figure 2.**
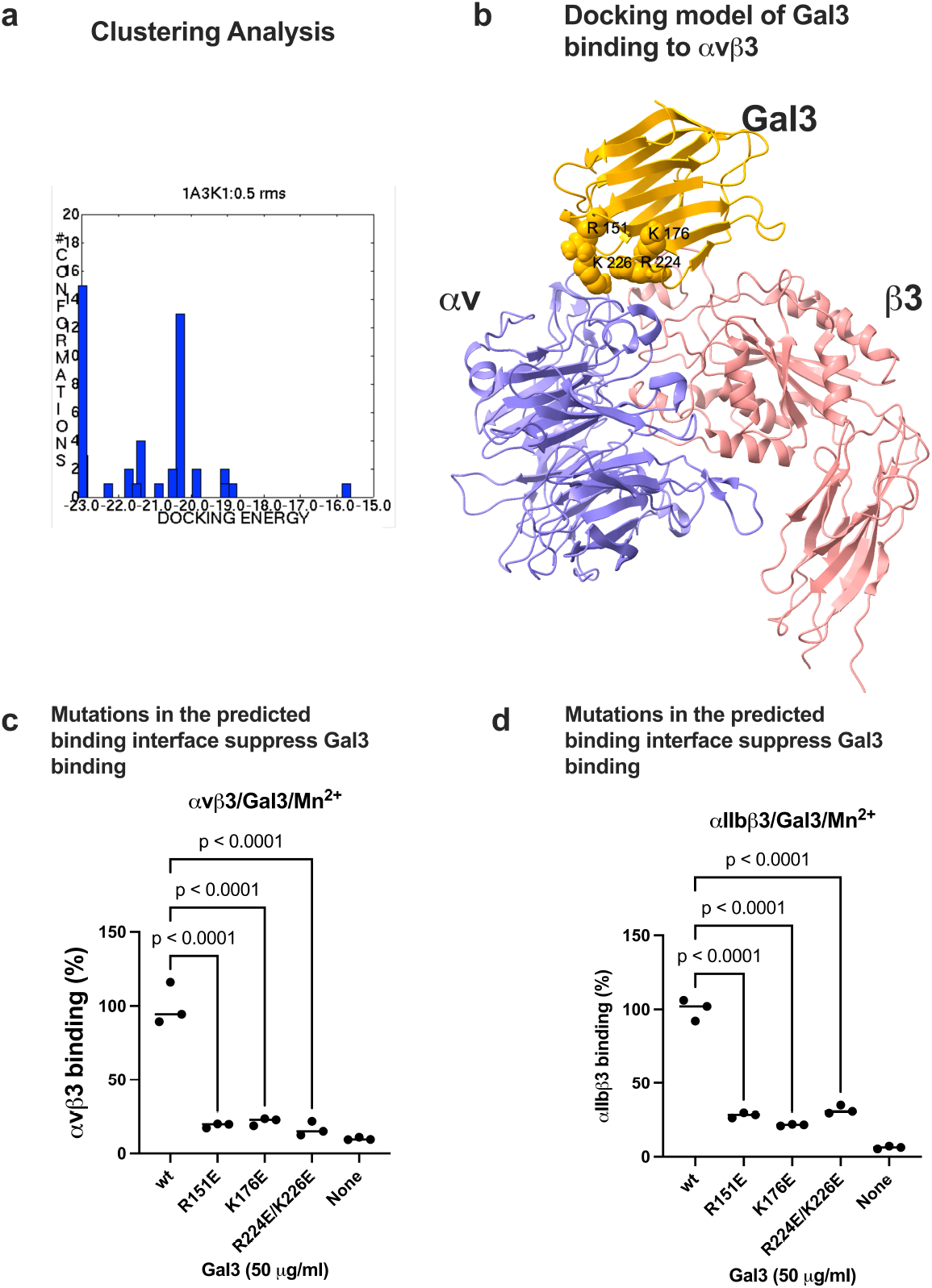
Docking simulation of Gal3 binding to αvβ3. Docking simulation between Gal3 and open headpiece αvβ3 (1L5G.pdb) was performed using autodock3 as described in the method section. (a) Clustering analysis of docked poses. The first cluster with docking energy -23 kcal/mol represent the most likely poses when Gal3 binds to site 1. (b) Docking model of Gal3 binding to site 1 of αvβ3 (1L5G.pdb). (c) Binding of Gal3 mutations to αvβ3 in 1 mM Mn^2+^. Point mutations of amino acid residues of Gal3 in the predicted binding interface effectively suppressed Gal3 binding to soluble αvβ3. (d) Binding of Gal3 mutations to αIIbβ3 in 1 mM Mn^2+^. Point mutations of amino acid residues of Gal3 in the predicted binding interface effectively suppressed Gal3 binding to soluble αIIbβ3. Data is shown as means +/-SD of triplicate experiments.

To prove if the docking model is correct, we introduced mutations of basic residues (Arg151, Lys176, and Arg224/Arg226) in the predicted integrin binding interface to Glu (charge reversal mutation). We found that R151E, K176E, and R224E/R226E mutations effectively suppressed Gal3 binding to integrins αvβ3 and αIIbβ3 in ELISA-type binding assays, suggesting that these amino acid residues are critical for binding to site 1 in these integrins. These findings are consistent with the docking model.

Several Gal3 residues are close to galactose in 3D structures of Gal3-galactose complex (His158, Asn160, Arg162, Val172, Cys173, Thr175, Asn174, Trp181, Gly182, Glu184, and Glu185) .(1A3K.pdb) are not in the Gal3-site 1 interface (Table 1), suggesting that galactose-Gal3 interaction may not be involved in integrin binding.

#### Gal3 binds to site 2 and induces allosteric integrin activation

Gal3 is a potent pro-inflammatory protein but the mechanism of this action is unclear. Previous studies showed that several pro-inflammatory cytokines (e.g., CX3CL1, CXCL12, CCL5, and CD40L) bind to site 2 of integrins and induce allosteric integrin activation and pro-inflammatory signaling (e.g., secretion of IL-6 and TNF) (19-22). Also, Gal3 enhances platelet aggregation and thrombosis (1). We thus hypothesized that Gal3 binds to site 2 and allosterically activates integrins, leading to pro-inflammatory signaling.

Docking simulation between Gal3 (1A3K.pdb) and integrin αvβ3 headpiece (inactive, closed headpiece form, 1JV2.pdb) predicts that Gal3 binds to the allosteric ligand binding site of αvβ3 (site 2) (Figure 3). The docking poses with the highest affinity (docking energy -21 kcal/mol) are clustered in the cluster 1, which represent the likely poses that binds to site 2.

**Figure 3.**
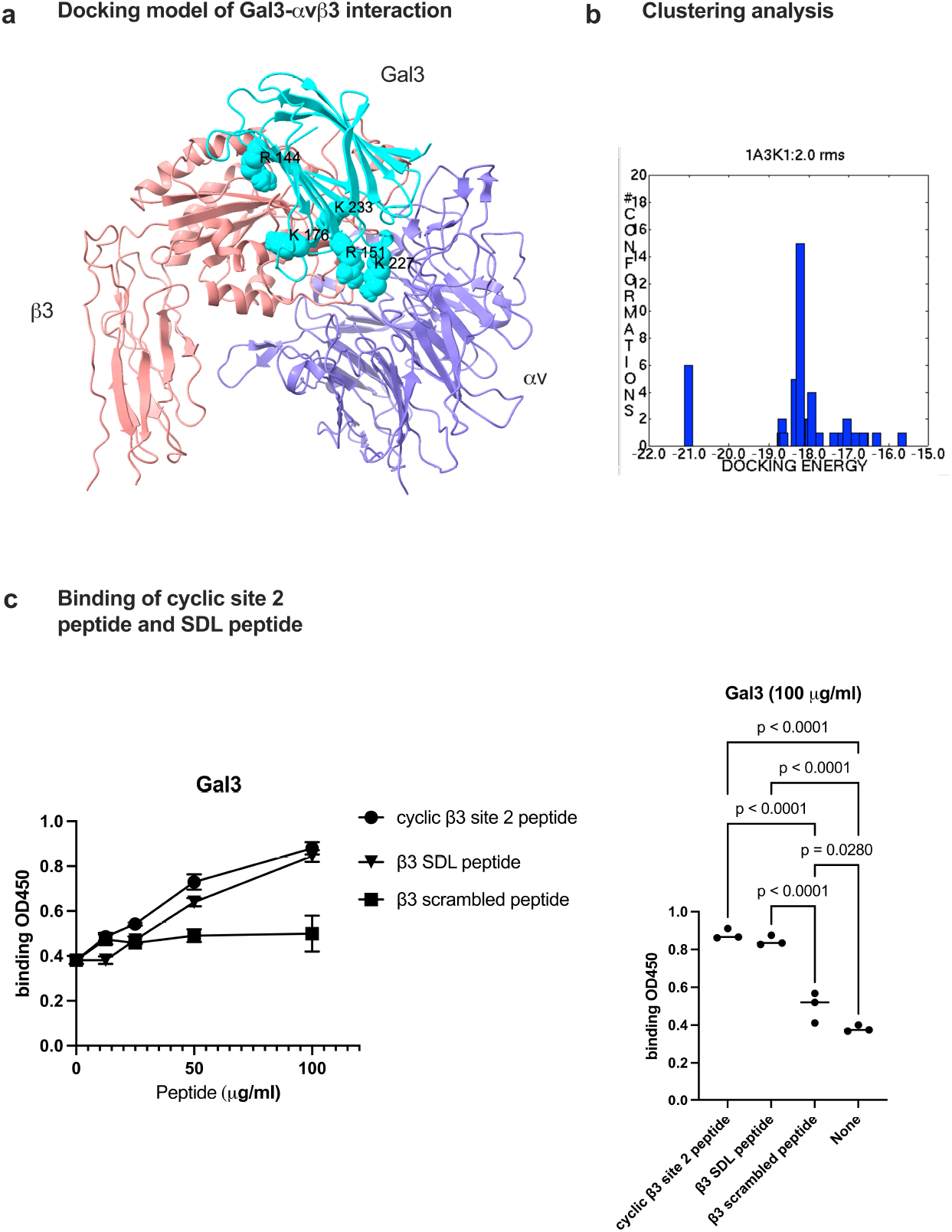
Docking simulation of galectin-3 to site 2 of αvβ3 (1JV2.pdb). Docking simulation between Gal3 and closed headpiece αvβ3 (1JV2) was performed using autodock3 as described in the method section. (a) The clustering analysis of docked poses. The first cluster with docking energy -21 kcal/mol represents the most likely poses when Gal3 binds to site 2. (b) The positions of amino acid residues in the predicted site 2 binding site. Amino acid residues involved in the interaction is shown in Table 2. (c) Binding of cyclic site 2 peptide and β3 SDL peptide to Gal3. Pepetide binding was performed as described in method section. Cyclic site 2 peptide and β3 SDL peptide and β3 SDL peptide bind significantly better to Gal3 than scramble peptide. Data is shown as means +/-SD of triplicate experiments.

Previous studies showed that ligands that bind to site 2 bind to site 2-peptide and/or SDL (specificity-determining loop)-peptide (Table 2) (2). We found that Gal3 binds to cyclic site 2 peptide and SDL peptides from β1 and β3 subunits (Figure 3). These findings suggest that Gal3 binds to site 2 and activates integrins.

**Table 2.** Predicted amino acid residues of Gal3 binding to site 2 of αvβ3.

| 1A3K (-21 kcal/mol) | $\alpha v$ (1JV2.pdb) | $\beta 3$ |
| --- | --- | --- |
| Leu114, Ile115, Val116, Pro117, Tyr118, Asn119, Leu120, Pro121, Pro123, Gly124, Gly125, **Arg144**, Ala146, Asp148, Gln150, **Arg151**, Gly152, Asn153, Asp154, Val155, His158, **Lys176**, Asn179, Trp181, **Lys227**, Asn229, Glu230, Ser232, **Lys233**, Leu234, Gly235, Ile236, Ser237, Ala245, Ser246, Tyr247, Thr248, | Glu15, Gly16, Lys42, Asn44, Val51, Glu52, His91, Arg122, Ser399 | Pro160, Met165, Ile167, Ser168, Glu171, Glu174, Asn175, Leu185, Pro186, Met187, Phe188, Lys191, His192, Val193, Lys209, Gln210, Ser211, <u>Cys273, His274, Val275, Gly276, Ser277, Asp278, Asn279, His280, Tyr281, Ser282, Ala283, Ser284, Thr285, Thr285</u> |

Amino acid residues within 0.6 nm between galectin-3 and $\alpha\beta 3$ were selected using Pdb Viewer (version 4.1). Amino acid residues of Gal3 selected for mutation are shown in bold. Amino acid residues in the site 2 peptide are underlined.

Binding of Gal3 to site 2 was studied in ELISA-type activation assays. Wells of 96-well microtiter plate were coated with fibrinogen fragments (γC399tr for αvβ3, and γC390-411 for αIIbβ3) and incubated with soluble αvβ3 or αIIbβ3 and bound integrins were quantified using anti-β3 mab and HRP-conjugated anti-mouse IgG. We found that Gal3 activated αvβ3 and αIIbβ3 in a dose-dependent manner (Figure 4).

**Figure 4.**
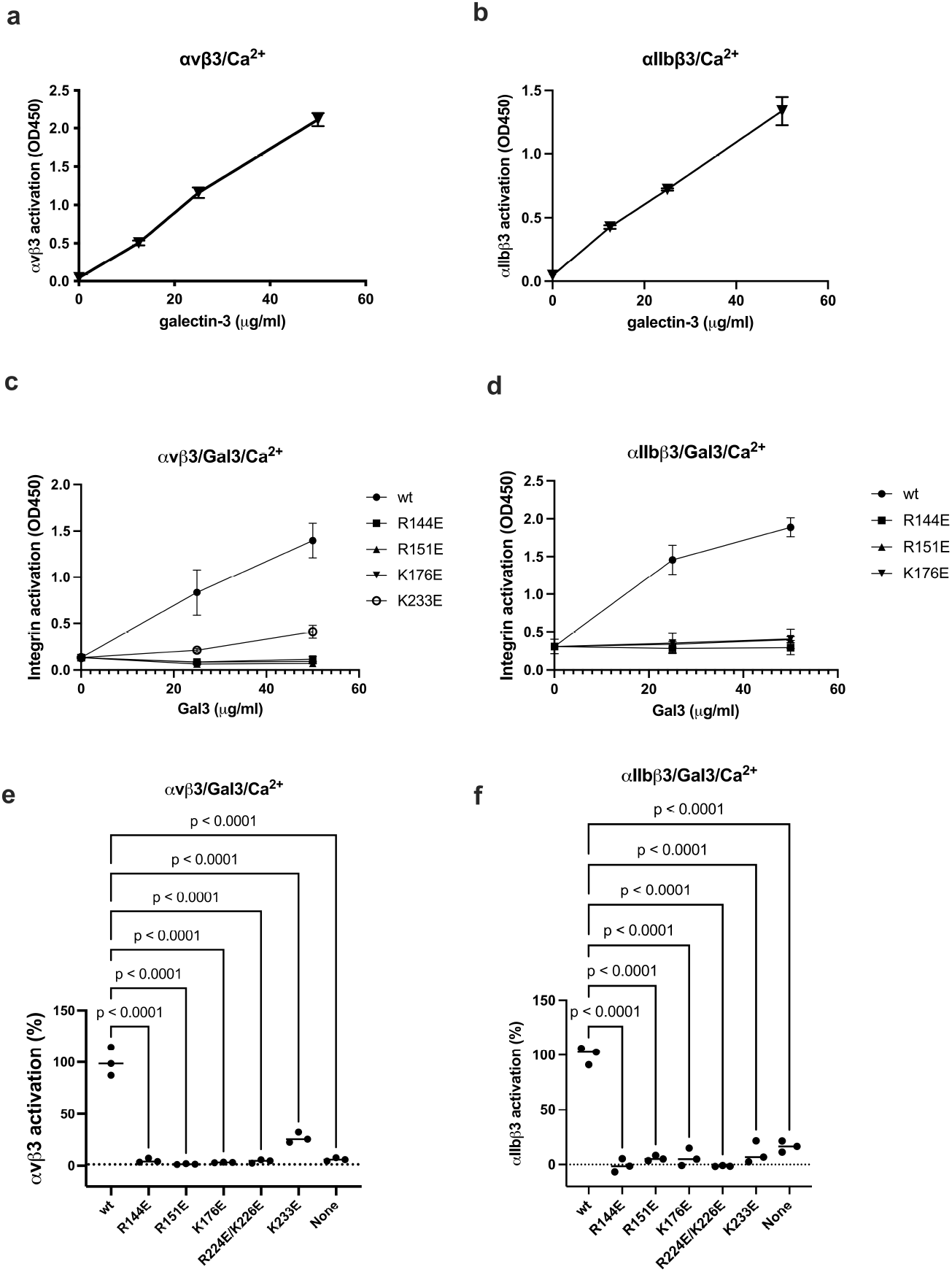
Gal3 binds to the allosteric binding site (site 2) and induce allosteric activation of soluble integrins. (a)(b) Activation of soluble integrins. Gal3 was incubated with soluble αvβ3 (a) or αIIbβ3 (b) in 1 mM Ca^2+^. Bound integrins were measured using anti-human β3 and HRP-conjugated anti-mouse IgG. (c-f). Mutations of the site 2 binding interface of Gal3 inhibit activation of soluble integrin αvβ3 (c) and soluble αIIbβ3 (d). (e and f) Integrin activation by Gal3 mutations (at 50 µg/ml). Data is shown as means +/-SD of triplicate experiments. (5-22-2024)

Amino acid residues involved in Gal3-αvβ3 interaction are shown in Table 2. We noticed that site 2-binding interface of Gal3 overlap with that of site 1-binding interface. We studied mutation of several basic residues (Arg144, Arg151, Lys176, Arg224/Lys226, Lys233) in the αvβ3-binding interface of Gal3 and mutated to Glu. These mutants effectively suppressed the ability of Gal3 to activate sol αvβ3 and αIIbβ3 (Figure 4), suggesting that these amino acid residues are involved in site 2 binding, which is consistent with the docking model.

### Gal3 binds to site 2 and induces allosteric activation of soluble α5β1 integrin

Previous studies showed that Gal3 binds to a5β1 purified transmembrane glycoprotein *α*5*β*1 integrin into a microcavity-suspended lipid bilayer (MSLB) platform of complex lipid composition in an activationg cation (Mn^2+^) (17). It is unclear if α5β1 is activated by Gal3 by binding to site 2. Docking simulation predicts that Gal3 binds to site 2 of α5β1 at comparable affinity to that of αvβ3 site 2. Predicted amino acid residues of Gal3 involved in α5β1 binding are shown in Table 3. We found that wt Gal3 activated soluble α5β1 (ectodomain) in 1 mM Ca^2+^in activation assays using biotinylated soluble α5β1 (Figure 5). Interestingly, several Gal3 mutants (R144E, R151E, K176E, and K227E) in the site 2-binding interface inhibited α5β1 activation by Gal3, suggesting that α5β1, αvβ3, and αIIbβ3 bind to site 2 in a similar manner. These findings suggest that Gal3-induced allosteric activation is not limited to β3 integrins.

**Table 3.**
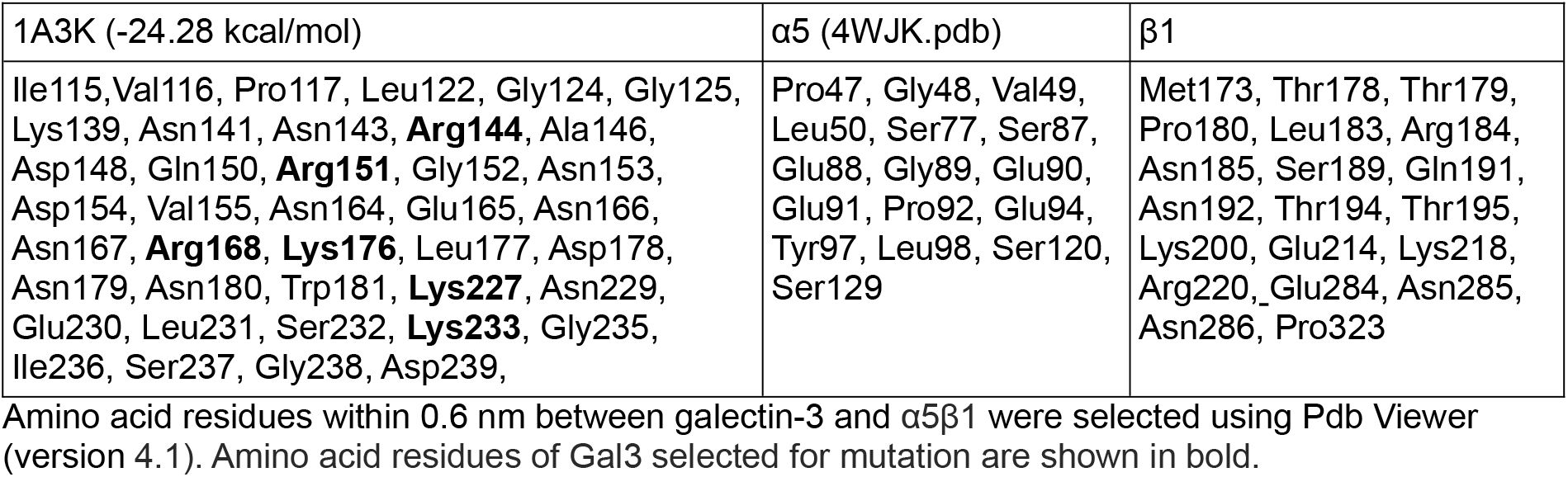
Predicted amino acid residues involved in Gal3 and α5β1 site2 binding.

**Figure 5.**
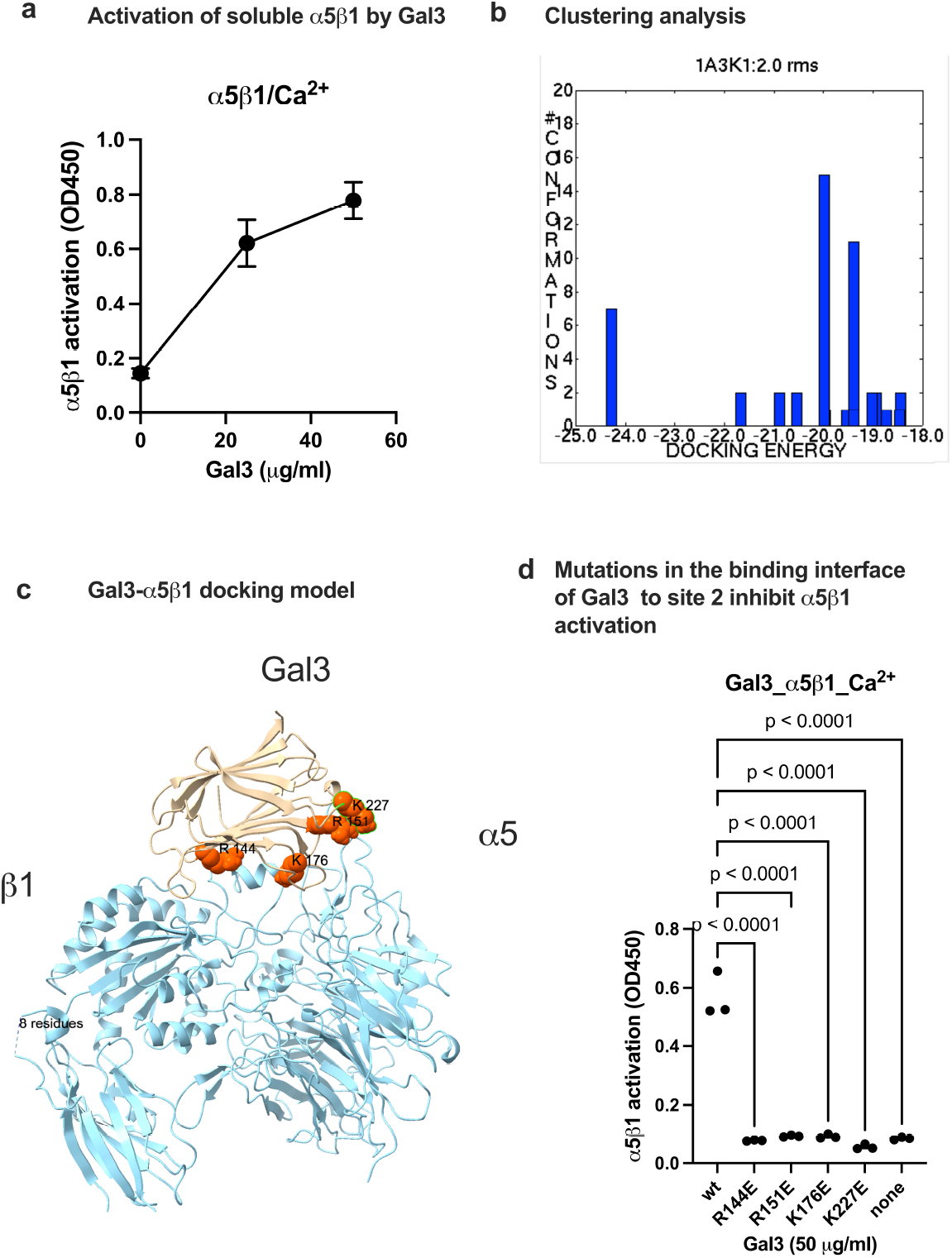
Gal3 activates α5β1 by binding to site 2. Docking simulation of α5β1-Gal3 interaction. Docking simulation of α5β1 (4wjk.pdb) and Gal3 (1A3K.pdb) was carried out as described in the Method section. a) Activation of soluble α5β1 by Gal3 in 1 mM Ca^2+^ in a deose-dependent manner. α5β1 activation was assayed as described for αvβ3 except that His-tagged soluble α5β1 was used. α5β1 was quantified using anti-His antibody conjugated with HRP. (b) Clustering analysis. The first cluster with docking energy -24.26 Kcal/mol represent the poses that bind to site 2 and activate α5β1. Docking simulation was performed as described in the method section. (c) Docking model of Gal3 binding to α5β1 in the first cluster. (d) Effect of mutations in the predicted site 2 binding interface of Gal3 to α5β1. Data is shown as means +/-SD of triplicate experiments.

### Anti-inflammatory FGF1, NRG1, and IVM inhibited Gal3-induced integrin activation

Previous study showed that anti-inflammatory FGF1 binds to site 2 but did not activate integrins. Instead FGF1 inhibited integrin activation by proinflammatory FGF2 (9). Also, anti-inflammatory NRG1 inhibited allosteric integrin activation induced by multiple pro-inflammatory cytokines (e.g., CCL5) (2). We also discovered that anti-inflammatory IVM, anti-parasite agent, binds to the center of site 2 and inhibited integrin activation induced by multiple inflammatory cytokines (e.g., TNF, CCL5, CXCL12) (8), suggesting that IVM is a site 2 antagonist. We hypothesized that IVM may inhibit integrin activation. We found that IVM, NRG1 and FGF1 (in this order) inhibited integrin activation induced by Gal-3 (Figure 6), suggesting that these anti-inflammatory agents act as antagonists for Gal-3 induced inflammatory signaling.

**Figure 6.**
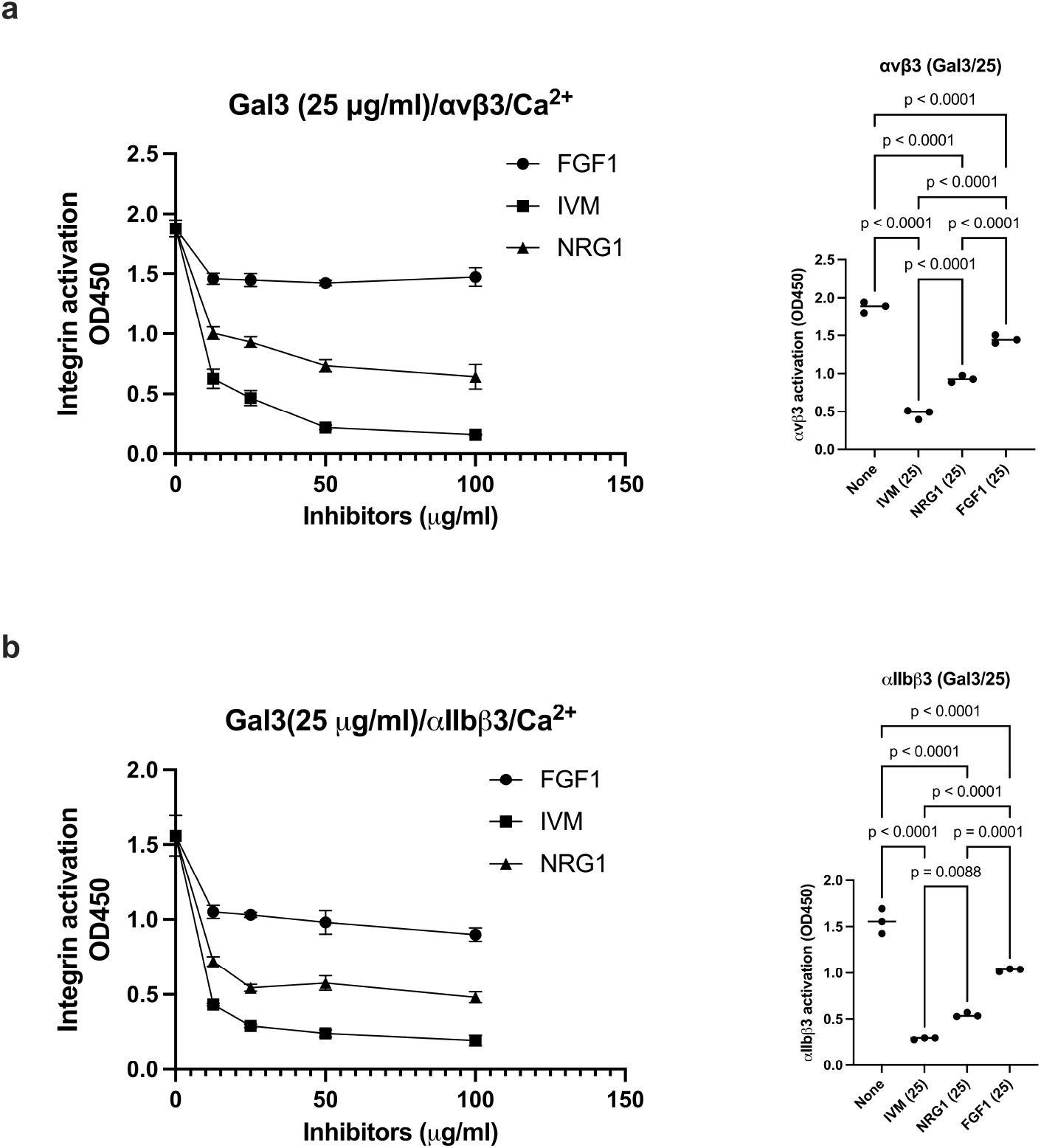
Anti-inflammatory FGF1, NRG1, and IVM suppress Gal3-induced integrin activation of αvβ3 (a) and αIIbβ3 (b). Activation of soluble αvβ3 (a) or αIIbβ3 (b) by Gal3 (25 µg/ml) in 1 mM Ca^2+^ was measured as described in methods section. The effect of FGF1, IVM, or NRG1 (up to 100 µg/ml) was studied. Data is shown as means +/-SD of triplicate experiments. The statistical analysis of the effect of FGF1, NRG1, and IVM at 25 µg/ml was performed.

Lactose did not affect Gal3-induced integrin activation, which is consistent with the docking model that lactose-binding site and site 2-binding site are distinct.

## Discussion

In the present study, we first establish that Gal3 binds to the classical RGD-binding site (site 1) of soluble integrin αvβ3 and αIIbβ3. RGDfV inhibited Gal3 binding to αvβ3 and eptifibatide inhibited Gal3 binding to αIIbβ3 in 1 mM Mn^2+^, suggesting that Gal3 binds to site 1 in an RGD-depedent manner.

Notably, this binding of Gal3 to integrins was not affected by lactose, suggesting that glycans are not involved in site 1 binding. Notably, the integrin-binding interface of Gal3 does not include known carbohydrate-binding site, which is consistent with the idea that Gal3-integrin binding is not dependent on glycan binding. Docking simulation predicts that Gal3 binds to site1 of αvβ3. Point mutations in the predicted site 1 binding interface of Gal3 effectively suppressed Gal3 binding to site 1 of soluble integrin αvβ3 and αIIbβ3, consistent with the docking model. Also, these findings suggest that Gal3 is similar to known integrin ligands. We found that Gal3 binds to soluble αIIbβ3 (Figure 1) in 1 mM Mn2+

### Since Gal3 is known to be one of the most pro-inflammatory proteins

Previous studies showed several inflammatory cytokines (e.g., CCL5, TNF, CX3CL1, CXCL12, sPLA2-IIA) bind to site 2 and allosterically activate integrins αvβ3 and αIIbβ3. Therefore, we hypothesized that Gal3 binds to site 2 and acts like inflammatory cytokines. Docking simulation predicts that Gal3 binds to site 2 of αvβ3, αIIbβ3 and a5b1. Consistently, Gal3 activated these integrins in 1 mM Ca^2+^. Point mutations in the predicted site-2 binding interface suppressed Gal3-induced activation of these integrins, suggesting that Gal3 binding to site 2 is required for Gal3-induced integrin activation. Gal3 bound to cyclic site 2 peptides and the SDL peptide from integrin β3 subunit, consistent with the idea that Gal3 binding to site 2 is required for Gal3-induced integrin activation. Notably, site 2-binding interfaces of Gal3 overlap when Gal3 binds to site 2 of three different integrins, suggesting that a single Gal3 antagonist to the site 2-binding site may effectively suppress pro-inflammatory action of Gal3. Gal3 is known to enhance platelet aggregation and thrombosis (1). The present study provides evidence that Gal3 binds to αIIbβ3 and this interaction may result in platelet activation and aggregation leading to pro-thrombotic action of Gal3.

Furthermore, we showed that FGF1 (9), ivermectin (8) and NRG1 (2) inhibited the Gal3-induced integrin activation, suggesting that these known anti-inflammatory cytokine or compound block Gal3 binding to site 2 as therapeutics for Gal3-induced pro-inflammatory and pro-thrombotic actions.

## Conclusion

In the present study, we performed docking simulation of interaction between Gal3 (CRD) and integrin αvβ3. The simulation predicted that Gal3 binds to site 1. We showed that Gal3 specifically binds to site 1 of soluble integrins αvβ3 and αIIbβ3 in cell-free conditions in 1 mM Mn^2+^. Lactose did not affect galactose-integrin binding, but integrin inhibitors did, suggesting that carbohydrate may not play a critical role in integrin binding. Mutations in the predicted binding interface to site 1 inhibited the binding. Furthermore, docking simulation predicted that Gal3 binds to site 2. Gal3 potently activated soluble β3 integrins in cell-free conditions in 1 mM Ca^2+^, suggesting that Gal3 induces integrin activation in an allosteric manner. Cyclic site 2-derived peptides bound to Gal3, which is consistent with the prediction that Gal3 binds to site 2. Point mutations in the predicted site 2-binding interface effectively suppressed Gal3-induced integrin activation. Notably, we showed that ivermectin, NRG1 and FGF1 inhibited integrin activation induced by Gal3, suggesting that Gal3 binding to site 2 is involved in integrin activation by Gal3. Allosteric activation of αIIbβ3 by Gal3 may trigger platelet aggregation and thrombosis. Allosteric activation of αvβ3 may trigger inflammatory signals trough site 2, and induce inflammatory signals. We propose that Gal3 binding to site 2 may be critically involved in pro-inflammatory and pro-thrombotic signals by Gal3, and that ivermectin, NRG1 and FGF1 have potential as antagonists for Gal3-mediated diseases.

## Materials and Methods

### Materials

Ivermectin (IVM) was obtained from eMolecules (San Diego, CA, USA). We used stock IVM (10 mM) in 95% ethanol and diluted it in water. Human soluble αvβ3 (IT3-H52E3) and αIIbβ3 (IT3-H52W8) were obtained from ThermoFischer (Waltham, MA, USA). Mab AV10 (to human β3) was kindly provided by Brunie Felding (Scripps Research Institute, La Jolla, CA, USA). HRP-conjugated anti-His tag (C-terminal) antibody was purchased from Qiagen (Valencia, CA, USA).

#### Galectin-3 (Gal3)

cDNA encoding the carbohydrate-binding domain of Gal3 (residues 114-250) [LIVPYNLPLPGGVVPRMLITILGTVKPNANRIALDFQRGNDVAFHFNPRFNENNRRVIVCNTKLDNNWGREERQSVFPFESGKPFKIQVLVEPDHFKVAVNDAHLLQYNHRVKKLNEISKLGISGDIDLTSASYTMI] were synthesized and subcloned into the BamHI/EcoRI site of pET28a vector. Protein synthesis was induced by IPTG in E. coli BL21. Protein was synthesized as insoluble inclusion bodies and purified in denaturing conditions (8 M urea) in Ni-NTA Sepharose and refolded as previously described (23).

#### Cyclic β3-site 2 peptide

We synthesized the 29 mer cyclic β3 site 2 peptide C260-RLAGIV[QPNDGSHVGSDNHYSASTTM]C288 (C273 is changed to S273) by inserting oligonucleotides encoding this sequence into the BamHI/EcoRI site of pGEX2T vector as described (2).

#### β3 SDL peptide

The cDNA fragment encoding the specificity-determining loop (SDL) (residues 158– 188 of β3 [DKPVSPYMYISPPEALENPCYDMKTTCLPMF]) was synthesized and subcloned into the Bam HI/Eco RI site of pGEX2T vector (2). Proteins were synthesized in E. coli BL21 and purified using Glutathione Sepharose affinity chromatography.

#### Docking simulation

Docking simulation between the C-terminal region of galectin-3 and integrin αvβ3 or α5β1 was performed as described above (24). The ligand was presently compiled to a maximum size of 1024 atoms. Atomic solvation parameters and fractional volumes were assigned to the protein atoms by using the AddSol utility, and grid maps were calculated by using the AutoGrid utility in AutoDock 3.05. A grid map with 127 × 127 × 127 points and a grid point spacing of 0.603 Å included the headpiece of αvβ3 or α5β1. Kollman ‘united-atom’ charges were used. AutoDock 3.05 uses a Lamarckian genetic algorithm (LGA) that couples a typical Darwinian genetic algorithm for global searching with the Solis and Wets algorithm for local searching. The LGA parameters were defined as follows: the initial population of random individuals had a size of 50 individuals; each docking was terminated with a maximum number of 1 × 106 energy evaluations or a maximum number of 27,000 generations, whichever came first; and mutation and crossover rates were set at 0.02 and 0.80, respectively. An elitism value of 1 was applied, which ensured that the top-ranked individual in the population always survived into the next generation. A maximum of 300 iterations per local search were used. The probability of performing a local search on an individual was 0.06, whereas the maximum number of consecutive successes or failures before doubling or halving the search step size was 4.

## Methods

### ELISA-type binding assay

Wells of 96 well microtiter plate were incubated with Gal3 in PBS for 1h at room temperature and the remaining binding sites were blocked by incubating with 0.1% heat-treated BSA (80°C for 10 min). Well were incubated with recombinant soluble αvβ3 or αIIbβ3 (1 ug/ml) in HEPES-Tyrodes buffer with 1 mM Mn^2+^ for 1 h at room temperature. After washing the wells, bound integrins were quantified using anti-β3 mAb (AV10) and HRP-conjugated anti-GST and peroxidase substrate.

### ELISA-type activation assay

Wells of 96 well microtiter plate were incubated with recombinant fibrinogen fragment γC399tr (specific ligand for αvβ3) or γC390-411 (specific to αIIbβ3) (50 µg/ml in PBS) for 1 h at room temperature. Remaining binding sites were blocked by incubating with 0.1% heat-treated BSA (80 C for 10 min). Well were incubated with recombinant soluble αvβ3 or αIIbβ3 (1 µg/ml) in the presence of Gal3 in HEPES-Tyrodes buffer with 1 mM Ca^2+^ for 1 h at room temperature. After washing unbound integrins, bound integrins were quantified using anti-β3 mAb (AV10) and HRP-conjugated anti-mIgG and peroxidase substrate.

### Site-directed mutagenesis

was performed using the QuickChange method (25).

### Statistical Analysis

We used Prism 10 (Graphpad Software, Boston, MA, USA) to test treatment differences using ANOVA and Tukey multiple comparison tests to control the global type I error.

## Data Availability Statement

We will share existing datasets or raw data that have been analyzed in the manuscript upon request.

## Author contribution

Conceptualization and writing—review and editing, Y.T.; data curation, Y.K.T. and C.-Y.W. All authors have read and agreed to the published version of the manuscript.

